# Collagen assembly and turnover imaged with a CRISPR-Cas9 engineered Dendra2 tag

**DOI:** 10.1101/331496

**Authors:** Adam Pickard, Antony Adamson, Yinhui Lu, Joan Chang, Richa Garva, Nigel Hodson, Karl E. Kadler

## Abstract

Electron microscopy has been the “gold standard” for studying collagen networks but dynamic information on how cells synthesise the networks has been lacking. Live imaging methods have been unable to distinguish newly-synthesised fibrils from pre-existing fibrils and intracellular collagen. Here, we tagged endogenous collagen-I using CRISPR-Cas9 with photoswitchable Dendra2 and demonstrate live cells synthesising, migrating on, and interacting with, collagen fibrils. This strategy is applicable for other long half-life proteins.

---

There are 28 genetically distinct collagens in vertebrates of which the most abundant is collagen-I ^1^. This collagen occurs as centimetre-long fibrils in the extracellular matrix and binds integrins^2^, making it essential for metazoan development^3^. The importance of the fibrils is exemplified in genetic diseases of collagen-I, for example the brittle bone disease osteogenesis imperfecta^4^, and in fibroproliferative diseases, often involving TGF-ß1 stimulated collagen synthesis^5^, in which excess or ectopic deposition causes organ failure, often with fatal consequences^6^. Although the fibrils are relatively long (>1 cm) their narrow diameters (typically 50 – 100 nm) limit the information that can be obtained by light and fluorescence microscopy methods. Therefore electron microscopy (predominately transmission EM and serial block face-scanning EM) has become the gold standard for studying collagen transport and fibril assembly (for example, see^7^). However, the major setbacks are the lack of dynamic data and the inability to distinguish fibrils that are newly deposited from those that are pre-existing and might be undergoing turnover.

Photoactivatable fluorescent proteins are capable of undergoing changes in fluorescent emissions in response to specific wavelengths of light irradiation. One such protein is Dendra2, which is a monomeric green-to-red photoconvertible fluorescent protein^8^ that has gained popularity in studying intracellular protein movements, aggregation and turnover (e.g. see ^8–11^) but has not previously been used to assess its usefulness in studying protein polymerisation on the scale of collagen fibrils in the extracellular matrix.

As collagen is a secreted extracellular matrix protein, we introduced sequences coding for Dendra2 3’ of the signal peptide of *Col1a2* to facilitate targeting of the modified polypeptide to the lumen of the endoplasmic reticulum. Collagen-I is synthesised as a precursor procollagen-I molecule containing two proa1(I) polypeptides and one proa2(I) polypeptide chain, encoded by *Colla1* and *Col1a2* genes, respectively (**Fig. 1a**). Proα2(I) differs in domain structure from proα1(I) in lacking a globular N-terminal domain. A minor variant of procollagen-1 is the [proα1(I)]_3_ homotrimer. We chose to modify the proα2(I) chain by CRISPR-Cas9 based tagging to specifically target the more abundant heterotrimeric isoform of collagen-I. We were careful to select an insertion site that would not disrupt collagen fibril formation. This required the identification of a site in the proa2(I) chain that would not disrupt the C-terminal propeptides (which contain the sites for chain selection^12^), disrupt removal of the C-propeptides (which is required for fibril formation^13^), remove the binding sites for fibril formation^14^ or alter the repeating Gly-X-Y structure of the collagen molecule^15^). We reasoned that the only possible site for insertion of Dendra2 was N-terminal of the N-propeptides, i.e. at the most N-terminus of the molecule (**Fig. 1b**). The N-propeptides can be proteolytically removed by ADAMTS 2, 3 and 14^16^, and, moreover, defects in removal of the N-propeptides are associated with soft tissue frailty in people with the Ehlers Danlos syndrome type VII^17^. Furthermore, studies have shown that the N-propeptides are not completely removed from all procollagen molecules ^18^ and collagen synthesis might be dysregulated by defective removal of the N-propeptide^19^. For these reasons, we opted not to engineer out the N-propeptide ADAMTS cleavage site. A further consideration was to tag endogenous proa2(I) to avoid possible complications caused by overexpression of collagen-I.

**Fig. 1:**
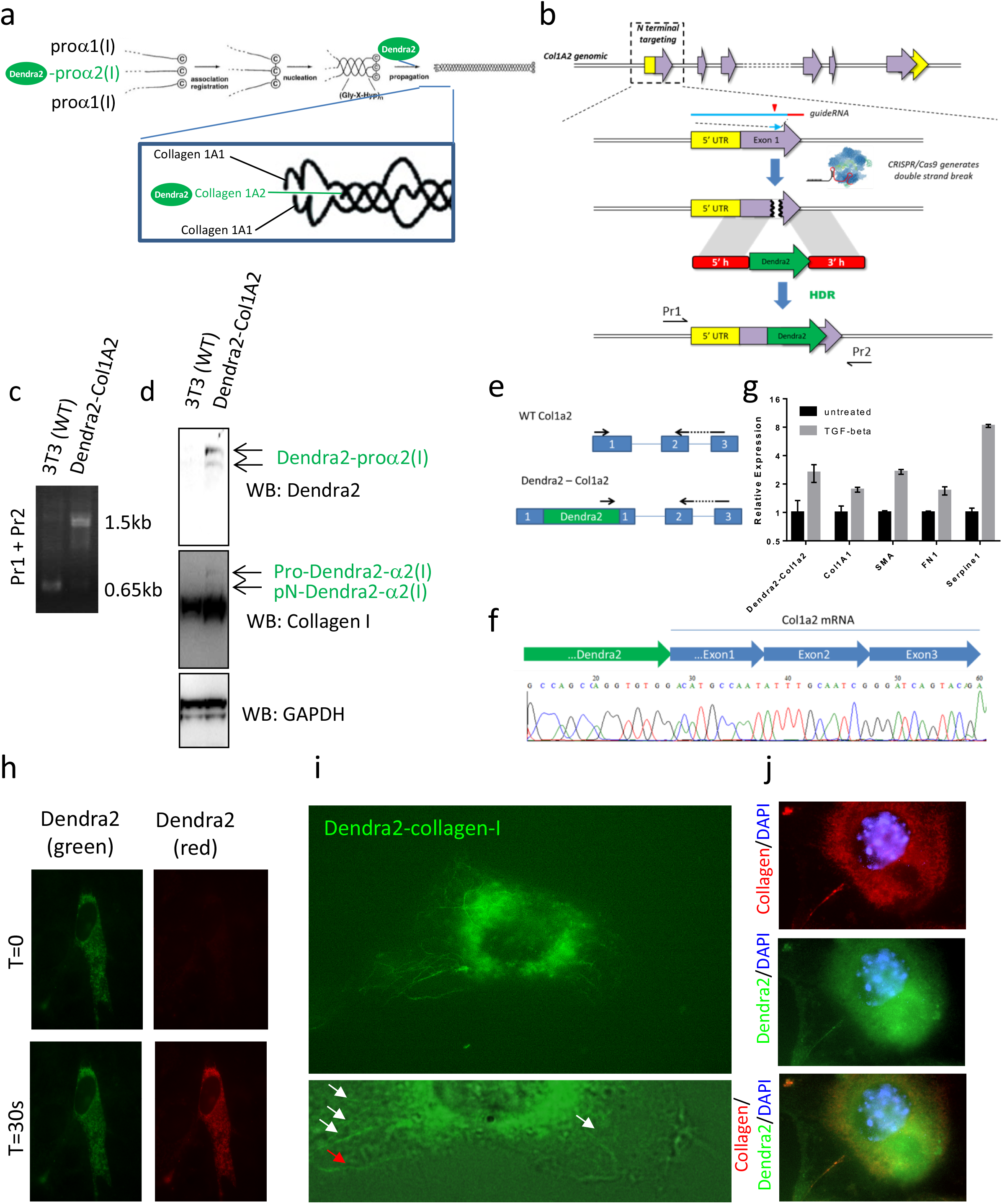
CRISPR-Cas9 mediated tagging of collagen-I containing fibres with Dendra2. (**a**) Integration of Dendra2 into the N-terminus of proa2(I). The short N-terminus of proa2(I) provides space within the N-propeptide for incorporation of Dendra2 into the collagen-I molecule (modified from ^27^). (**b**) Integration of sequences encoding Dendra2 into the *Col1a2* gene used a guide RNA immediately downstream of the signal peptide, directing Cas9 to exon 1 of *Col1a2*. A repair template encoding the Dendra2 coding sequence flanked by 800 bp homolgy arms. Primers to validate intergation of the repair template are shown. (**c**) PCR validation of Dendra2 integraton to the *Col1a2* gene. DNA extracted from a single cell clone identified as Dendra2 positive by FACS. Primers are as shown in (b). (**d**) Western blot validation of Dendra2 knock-in in the same single cell clone compared with un-edited NIH3T3. (**e**) Real-time PCR primers were designed to detect editing. PCR products from edited cells were validated by sanger sequencing (**f**). (**g**) The editted *Dendra2-Col1a2* gene retrained responsiveness to TGF-βl treatment and was induced transcriptionally following 48 hours treatment. (**h**) Photoswitching of Dendra2-Col1a2 NIH3T3 using 30 s exposure to 400 nm light. Images before and after switch are shown. (**i**) Treatment of Dendra2-Col1a2 NIH3T3 cells with 200 μg/mL L-ascorbic acid for 5 days. Dendra positive fibrils are observed within the boundaries of the cell (white arrows) and also at the boundary of the cell (red arrow). (**j**) Indirect immunofluorescent detection of Dendra2 and collagen-I confirmed that the Dendra2 positive fibrils are collagen-I.

As proof-of principle for the tagging approach we generated single cell clones of NIH3T3 fibroblasts that express Dendra2-Col1a2 mRNA and protein (**Fig. 1c, d**). Sequencing of genomic DNA and cDNA generated from transcripts expressed from the Dendra2-Col1a2 locus verified the correct location of Denda2 (**Fig. 1e, f**). The expected induction of *Col1a2* transcription by pro-fibrotic stimuli was observed in these cells (**Fig. 1g**), indicative that Dendra2 tagging does not affect physiological responses to stimuli. Dendra2 maintained its photoswitchable properties when fused to proa2(I) (**Fig. 1h**). Super-resolution microscopy of live cells showed collagen fibril formation with addition of L-ascorbic acid (**Supplementary video 1 and 2, Fig. 1i**), and fibres formed on the underside of the cell (**Supplementary video 3**) suggesting that fibril assembly is a cell-controlled process occuring at the plasma membrane (**Fig. 1i**).

Dendra2-collagen-I fibrils were photoswitchable (**Supplementary Fig. 1**) and readily detected by antibodies that recognise native triple helical collagen (**Fig. 1j**). The ability of Dendra2-collagen-I to assemble into D-periodic fibrils was confirmed by EM and correlative AFM (**Fig. 2 a-c**). The *D*-periodicity was 65 ± 7.4 nm (**Fig. 2d** and **Supplementary Fig. 2**), and therefore within the expected range for collagen-I fibrils ^20^. We noted that the ends of fibrils were spiralled or looped, which is a feature not previously seen of collagen fibrils (**Fig. 2e**).

**Fig. 2:**
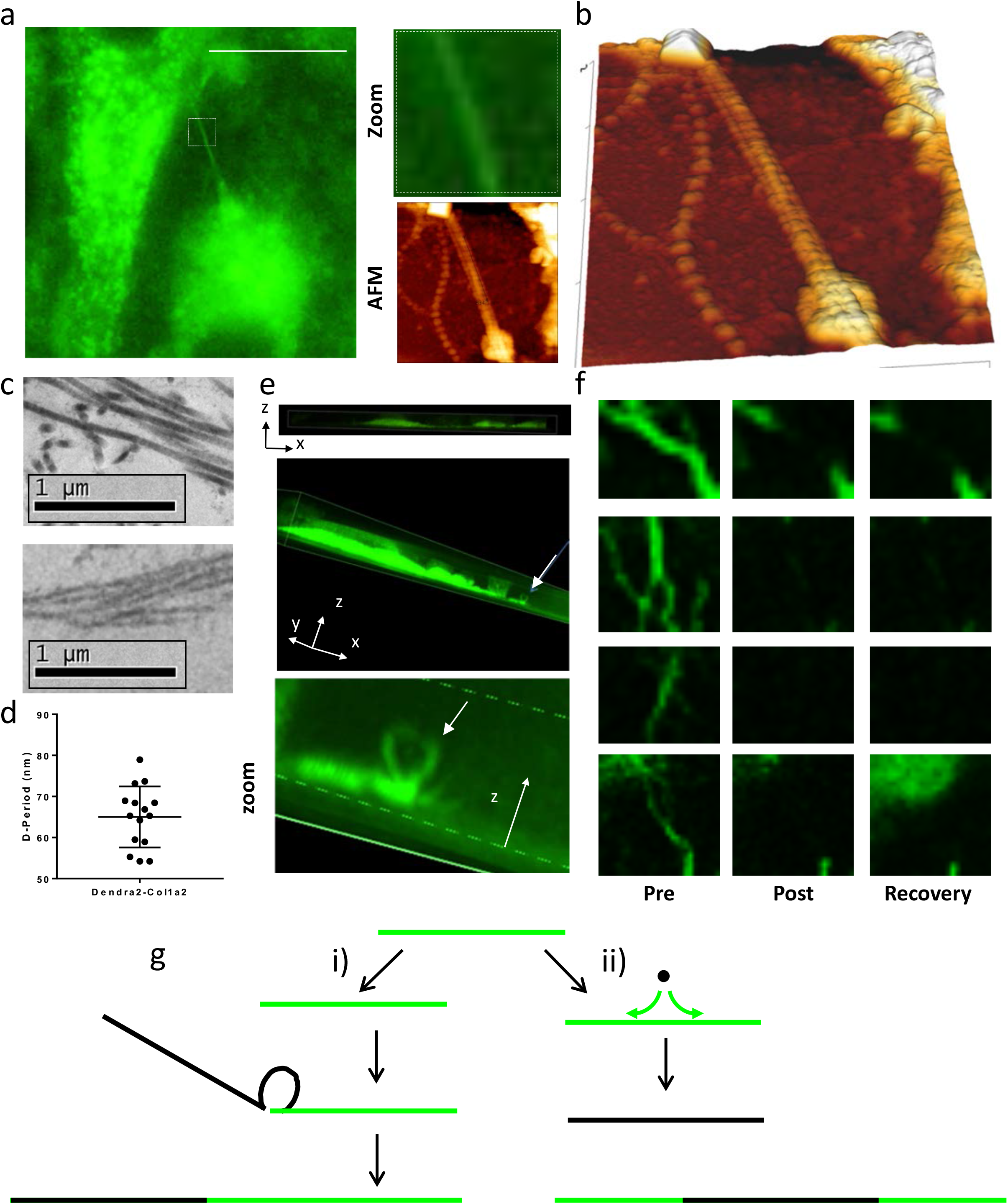
Collagen-I containing Dendra2-tagged pNα2(I) chain can form banded collagen fibrils. (**a**) Dendra2-Col1a2 editted NIH3T3 cells were grown for 5 days in L-ascorbic acid on Grid 500 μ-DISH (iBidi), this same region was then imaged using atomic force microscopy. (**b**) High resolution AFM identified multiple collagen fibrils within the same region of fluroescence signal. D-periodic banding was observed. (**c**) Transmission electron microscopy of collagen fibrils produced by uneditted (upper) and Dendra2-Col1a2 editted MC3T3 cells (lower). Fibrils are banded in both cultures. Scale bar, 1 μm. (**d**) Analysis of Dendra2-collagen-I fibrils by AFM (see also **Supplementary Fig. 2**) identified a mean *D*-periodicity of 65 ± 7.4 nm. (**e**) Z-stacked images of Dendra2-Col1a2 editted NIH3T3 cells showed fibrils on the underside of the cells and identified the prescence of looped structures at the fibril tips. (**f**) Photobleaching of Dendra2-collagen-I fibrils asscoiated with a single cell. Four different positions were chosen to bleach; each position showed no recovery of fluorescence over 24 hours. Different positions within the cell, and along the fibre were chosen (see also **Supplementary Fig. 3**). (**g**) A schematic of collagen fibril lengthening. Two scenarios are shown: scenario (i) shows fibril elongation by fusion of fibril ends to form longer fibrils; scenario (ii) shows fibril elongation by addition of collagen molecules to the shaft. No dynamics of collagen exchange or exchange in the shaft of an established fibre were observed. The evidence from photoswitching suggests scenario (i) is preferred.

We generated Dendra2-col1a2 in additional cell lines including MC3T3, a pre-osteoblast cell line, and visualised how these cells transport collagen fibres (**Supplementary Fig. 3 and Supplementary video 5**). Monitoring fluorescence recovery after photobleaching demonstrated that fibres are stable for at least 24 hours (**Fig. 2f, Supplementary Fig. 4**); there was no recovery of fluorescence and no degradation. In a similar approach, Dendra2-collagen-I fibres were photoswitched whilst on a micropattern of fibronectin, allowing single cells to be cultured. This approach confirmed that individual cells can produce a collagen matrix that is stable for at least 7 days in culture (**Supplementary Fig. 5**).

The fact that collagen fibril formation occurs in association with the plasma membrane raised the possibility that the length of collagen fibrils could be limited to the size of the cell. However, collagen fibrils *in vivo* can extend millimetres in length and therefore far longer than the length of individual fibroblasts^21^. Electron microscopy images have been interpreted to suggest that the occurrence of long fibrils is a result of end-to-end fusion^22^, but no definitive evidence has been available. Here we show that cells migrating over collagen fibrils pull and align the fibrils to facilitate end-to-end joining, thereby leading to rapid growth in fibril length (**Supplementary Fig. 3, Supplementary video 4** and schematic shown in **Fig. 2g**). It is possible that cells pull on specialised structures at the ends of fibrils, e.g. loops or spirals.

The use of a photoswitchable tag has also identified that collagen may have alternative routes of secretion (**Supplementary video 6**). We have observed that photoswitched (‘red’) collagen is removed from cells without passing through the Golgi, however newly synthesized collagen (‘green’) is transported through the Golgi. This suggests that collagen is either held within a post-Golgi compartment or is secreted via a Golgi-independent route. Whilst collagen can be secreted ^23^, we observe a pool of collagen that can remain within the Golgi for 2-3 hours.

We have used Dendra2-collagen-I expressing cells to explore matrix turnover. The results showed that degradation required direct contact of cells with the fibrils and often involves kinking of the collagen fibril before the fluorescence signal is lost (**Supplementary video 7**). X-ray structure studies of collagen fibrils have suggested that the 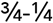 matrix metalloproteinase cleavage site in collagen-I^24^ is buried within the fibril structure and thereby inaccessible to these proteinases^25^. Kinking of the fibril might therefore be necessary to allow access of matrix metalloproteinases to the 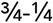 cleavage site.

The use of CRISPR-Cas9 to tag collagen combined with super-resolution microscopy has allowed visualisation of collagen fibril formation by living cells. The results show that collagen fibril assembly is a cell-controlled process that occurs at the cell surface. It was known that purified collagen-I can self-assemble into fibrils *in vitro* in the absence of cells^26^; however we observed no evidence of cell-free collagen fibril assembly. As assembly occurs on the underside of the cell, it is plausible that the cell creates a niche allowing the critical concentration of collagen to be reached to promote collagen fibril formation^13^. This hypothesis is supported by our observation that once fibril assembly begins the process is completed swiftly (**Supplementary video 2**). It is reasonable to consider that this approach could be applied to monitor the deposition of other fibrillar molecules. However, care should be taken to position the fluorescent tag within the target protein so that the fibril assembly sites are not masked. As the assembly and degradation of collagen both appear to occur at the cell surface, it is now tempting to suggest that diseases caused by adherent matrix assembly and degradation could be treated by targeting cell-collagen interactions.

---

**Supplementary Video 1: Generation of Dendra2-collagen-I positive fibrils by editted NIH3T3 cells**. Live cell imaging over 5 days on a Zeiss LSM 880 microscope with Airyscan mode. Cells were maintained at 37 °C with 5% CO2 and imaged every 30 minutes for four days.

**Supplementary Video 2: Generation of Dendra2-Col1a2 positive fibrils by editted NIH3T3 cells Title**. A region of Supplementary video 1 showing formation of collagen fibres.

**Supplementary Video 3: Confocal z-stack animated to shown collagen fibres underneath cells**. Z-stacks of Dendra2-Col1a2 editted NIH3T3 cell grown in L-ascorbic acid for 5 days on iBidi μ-dish. Multiple fibres are observed within the boundaries of a single cell. These fibres are also looped up away from the surface of the dish.

**Supplementary Video 4: Collagen fibrils can be joined by cells**. Dendra2-Col1a2 NIH3T3 cells were grown in L-ascorbic acid containing medium for 5 days and imaged every 30 minutes. A cell is seen pulling one fibril and joining this to a neighbouring fibril.

**Supplementary Video 5: Dendra2-Col1a2 editted MC3T3 cells produce collagen fibrils**. Dendra2-Col1a2 editted MC3T3 were grown in L-ascorbic acid supplemented medium for 24 hours and then imaged every 30 minutes for 4 days.

**Supplementary Video 6: Intracellular collagen turnover over 2 days**. A Dendra2-Col1a2 editted NIH3T3 cell was photoswitched with 30 s exposure to 400 nm light. Cells were imaged every 20 minutes for 2 days. The disappearance of photoswitched collagen occurs prior to generation of new ‘green’ Dendra-proα2(I). Timestamp is in hours.

**Supplementary Video 7: Cells are required to break collagen fibres**. Dendra2-Col1a2 editted NIH3T3 cells were used to produce a matrix containing Dendra2-collagen fibrils. Cultures were then decellularised and treated with DNase1. The matrix was pohotoswitched and then re-seeded with immortalised tail tendon fibroblasts. Images were taken every 30 minutes over 24 hours.

## Methods

See online Supplementary materials for further details.

### Cell culture

Immortalised mouse embryonic fibroblasts, NIH3T3, were maintained in DMEM supplemented with 10% newborn bovine calf serum and penicillin and streptomycin. The preosteoblast cell line MC3T3 were maintained in alpha modification MEM supplemented with 10% fetal bovine serum and penicillin and streptomycin. For imaging, cells were grown on ibitreat μ-Dish (Ibidi, Germany). Where indicated NIH3T3 and MC3T3 cell lines were cultured with 20 and 200 μg/mL L-ascorbic acid, respectively, to induce collagen fibre formation. Immortalised tail tendon fibroblasts were generated as previously described and were maintained in DMEM/F12 1:1 mixture supplemented with 10% FBS, penicillin and streptomycin. Cell survival was assessed using Alamar blue.

### CRISPR-Cas9 Knock in design

We used CRISPR-Cas9 to generate the Col1A2-Dendra2 knock in cell line. Using the Sanger CRISPR design webtool (http://www.sanger.ac.uk/htgt/wge/) we selected a guide RNA (*ACTTACATTGGCATGTTGCT* <u>**AGG**</u>) targeting exon 1 of the genomic sequence in order to generate a double strand break immediately after the sequence encoding the signal peptide. The guide was delivered as RNA oligos (Integrated DNA Technologies, Coralville, US). A double stranded DNA repair template was assembled by Gibson Assembly (NEB) using the primers in Table 1. Briefly, the 5’ and 3’ homology arms (800 bp each) were amplified from mouse genomic DNA. The Dendra2 mouse codon optimised coding sequence with a flexible linker sequence was synthesised (Genscript, US).

**Table 1:**
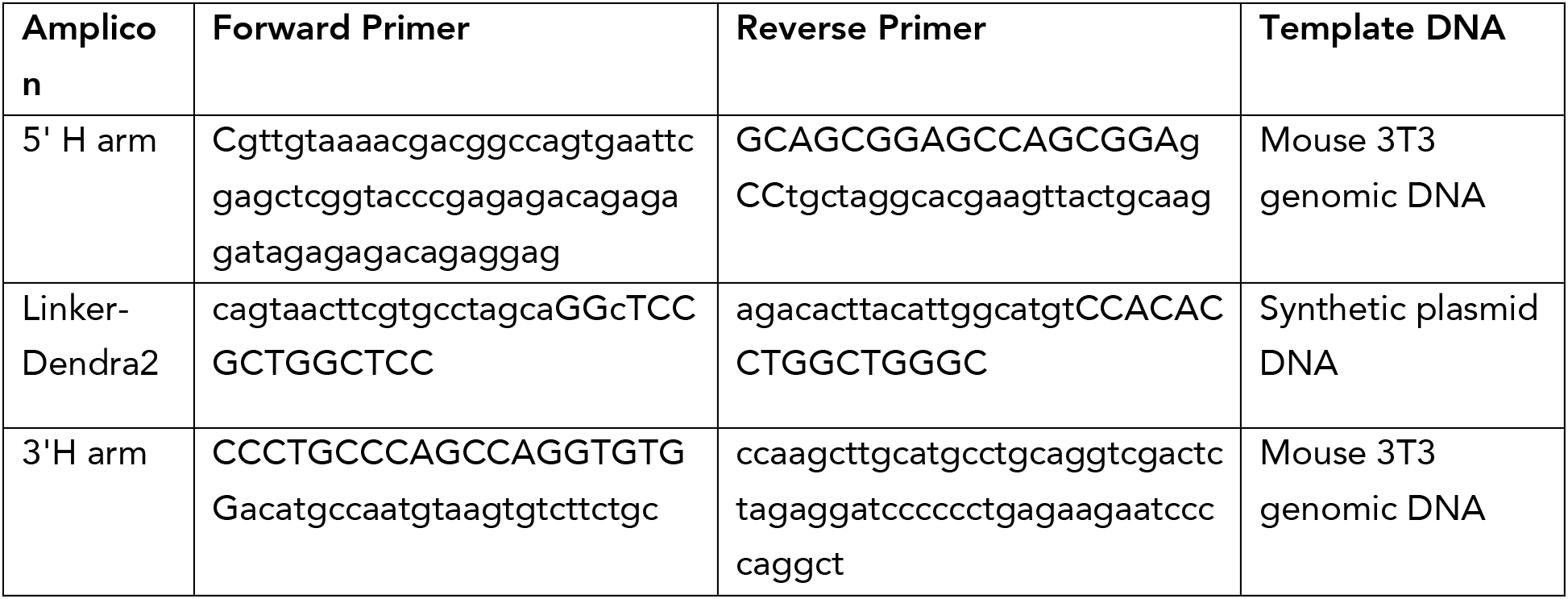
Primers used to generate repair template.

### CRISPR-Cas9 Delivery

Cells were seeded 24 hours prior to transfection of the repair template, using Fugene 6 (Promega) at a ratio of 3 μL per 1 μg DNA, cells were maintained in transfection mixture overnight and then fresh medium was added for approximately 6 hours. Cells were then transfected overnight with 100 pmol of tracrRNA and crRNA complexed with 4 pmol recombinant Cas9 protein using RNAiMAX (Thermo Fisher). Medium was then replaced and cells were grown for 48 hours.

### CRISPR-Cas9 Validation

Dendra2 positive populations were sorted by FACS and grown to sufficient numbers for validation of editing. Cells then sorted into single cells clones. All assays were performed on single cell sorted populations. Individual cells were then grown to sufficient numbers for validation. Primers for validation of the knockin (PR1 and PR2) were as follows: F-GGCAAGGGCGAGAGAGG and R-TTTTCTCCGACAGATTAGAGGGC. Real time validation of dendra2-col1a2 transcripts were assessed in single cell clones treated with 2.5 ng/mL TGF-ß1 for 48 hours using the primers indicated in Table 2, real time PCR was normalised to the geometric mean of Rplp0, Gapdh and Actb. The sequence of the Dendra2-col1a2 transcript was then validated by Sanger sequencing. Western blotting was performed on 4-12% tris-glycine gels with 25 μg cell lysate. Antibodies used in this study were Collagen (Gentaur, OARA02579, dilution 1:2000), Dendra2 (Origene, TA180094, dilution 1:500) and GAPDH (Sigma, G8795, dilution 1:10,000).

**Table 2:**
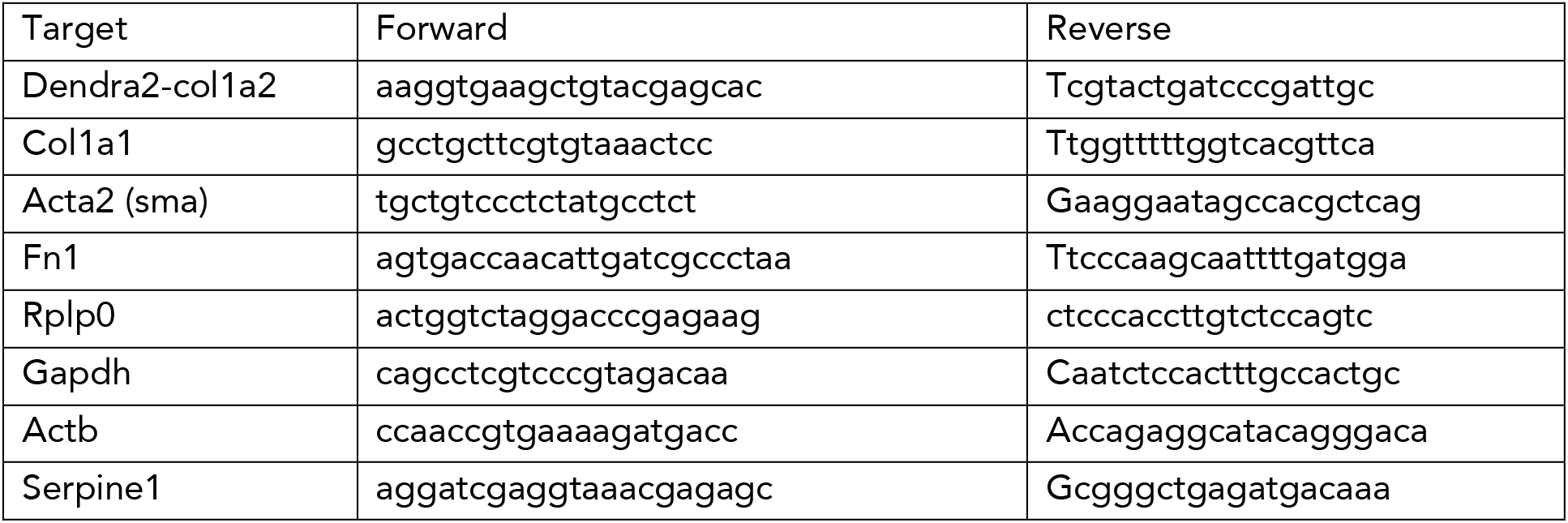
Primers used for real time PCR

### Imaging

Fig. 1h and i, 2c, 2h-j, Supplementary Fig. 1, 2 and video 4, 6 and 7 were acquired on an Eclipse Ti inverted microscope (Nikon) using a 60x objective, the Nikon filter sets for GFP and mCherry and LED (Lumencor) fluorescent light sources each with 300 ms exposure. Photoswitching was performed using a 30 s exposure to UV LED light source (400 nm). This system was also used to generate fibronectin micropatterns, briefly patterns were generated using the Primo patterning module (Alveole) according to manufacturer’s protocols before coating with 10 μg/mL human plasma fibronectin (BD biosciences). Imaging software NIS Elements AR.46.00.0 and point visiting was used to allow multiple positions to be imaged within the same time-course and cells were maintained at 37°C and 5% CO2. The images were collected using a Retiga R6 (Q-Imaging) camera. Fig. 1j. Images were collected on a Zeiss Axioimager.D2 upright microscope using a 100x objective and captured using a Coolsnap HQ2 camera (Photometrics) through Micromanager software v1.4.23. Following 4% PFA fixation cells were permeabilised and stained with antibodies to Dendra2 (dilution 1:100) and collagen (dilution 1:200), secondary antibodies used were goat-anti-mouse488 (Cell Signaling; 4408, dilution 1:400) and goat anti-rabbit-Cy5 (Invitrogen; A10523, dilution 1:400). Specific band pass filter sets for FITC and Cy5 were used to prevent bleed through from one channel to the next. All images were processed and analysed using Fiji ImageJ (http://imagej.net/Fiji/Downloads). Supplmental video 3, Fig. 2e were acquired on Nikon iSIM confocal microscope (40x magnification, 300 ms exposure). Fig. 2g was acquired using the Nikon A1R confocal microscope (60x magnification, 300 ms exposure). Supplementary video 1, 2 and 5 were acquired with a Fluar 20X 0.75NA objective at 1024 × 1024 resolution using a Zeiss LSM880 equipped with Airyscan detector set to super-resolution mode. Green fluorescence was excited at 488 nm and collected through a 495-550 nm filter. The 32 phase images were recombined using the airyscan processing tool in the Zeiss Zen 2 software and the image brightness and contrast adjusted using BestFit.

### Electron microscopy

Dendra2-Col1a2 edited MC3T3 were grown for 7 days in the presence of 200 μg/mL L-ascorbic acid, replenishing medium and ascorbic acid every 2 days. Following fixation in 2.5% glutaraldehyde/100 mM phosphate buffer (pH 7.2) for 30 minutes at RT cells and matrix were scraped from the culture dish. After at least 2 hours at 4 °C the samples were washed in 100 mM phosphate buffer (pH 7.2). After immersion in 2% osmium, 1.5% potassium ferro-cyanide in cacodylate buffer (pH 7.2) for one hour at room temperature, samples were then extensively washed in ddH2O, and fixed in 1% tannic acid in 0.1 M cacodylate buffer (pH 7.2) for 2 hours at 4 °C. Samples were thoroughly washed in ddH2O and incubated in 2% osmium tetroxide (in ddH2O) for 40 min at room temperature, before washing again in ddH2O at room temperature. This is followed by a final incubation step at 4 °C in 1% uranyl acetate (aqueous) overnight. Samples were then washed before infiltrated with a series of propylene oxide and TAAB 812 resin kit mix, with increasing resin concentration (2 hours in 30% resin, 2 hours in 50% resin, 2 hours in 75% resin, 3 × 1 hour in 100% resin). Samples are then embedded in capsules and cured at 60 °C for 12 hours. Thin sections (80 nm) were cut from the sample blocks and examined using the Tecnai 12 BioTwin electron microscope.

### Atomic force microscopy

Atomic force microscopy was performed using a JPK NanoWizard IV (JPK Instruments AG, Berlin, Germany) mounted on a Zeiss AX10 (Carl Zeiss Microscopy GmbH, Jena, Germany) inverted light microscope operating under JPK NanoWizard Control software (V 6.1.65). Images were captured using NuSense Scout 350R cantilevers (NuNano, Bristol, UK) with nominal spring constant, frequency and tip radius of 42 N/m, 350 kHz and < 10nm respectively. Height data were processed using JPK Data Processing software (V 6.1.65), and was 1^st^ order flattened prior to analysis.

### Decellularization

Decellularisation and recellularisation experiments were modified from ^28^. Briefly, Dendra2-Col1a2 NIH3T3 cells were cultured in ascorbic acid-containing medium for 5 days, before media was aspirated and cells washed gently with PBS. Pre-warmed extraction buffer (20 mM NH4OH, 0.5% Triton X-100 in PBS) was then added to plates and left for 2 minutes at 37 °C until no intact cell is visible under light microscope. The cell extract was aspirated and remaining matrix washed twice with PBS containing calcium and magnesium, followed by 10 μg/ml DNase I treatment at 37 °C for 30 minutes. Matrix was then washed twice with PBS containing calcium and magnesium, before recellularisation with immortilised tail tendon fibroblasts.

## Acknowledgements

Special thanks go to Peter March and David Spiller for help with long term imaging, to Gina Gamble and Kate Lewis from Nikon UK Limited for imaging help with the A1R and iSIM confocal microscopy systems, to Michael Jackson for help with single cell sorting, and to the staff in the Transgenic Unit for help with CRISPR-Cas9. The research was funded by Wellcome Trust Investigator and Wellcome Centre Core Awards to K.E.K. (110126/Z/15/Z and 203128/Z/16/Z). B.C. is funded by a Wellcome 4-year PhD studentship (210062/Z/17/Z). Light microscopes in the Bioimaging Facility were additionally supported by the University of Manchester Strategic Fund.

